# Annexin 5 is a key player in orchestrating the pollen sequestrome

**DOI:** 10.64898/2026.10.22.760708

**Authors:** Małgorzata Lichocka, Angelika Fiodor, Olga Iwańska, Anna Golisz-Mocydlarz, Monika Zakrzewska-Płaczek, Magdalena Górecka, Marta Grech-Baran, Szymon Kubala, Anna Anielska-Mazur, Norbert Lewkowicz, Izabela Barymow-Filoniuk, Agata L Starosta, Jacek Hennig, Magdalena Krzymowska

## Abstract

Like dry seeds, mature pollen grains store a large reservoir of RNA species that enables rapid activation and germination under favorable conditions. The molecular mechanisms governing the accumulation of these RNA reservoirs remain poorly understood. Here, we show that the Arabidopsis pollen-specific protein annexin 5 (ANN5) localizes to the nucleolus and exerts dose-dependent control over ribosome biogenesis, as evidenced by altered steady-state rRNA levels in mature pollen. Ectopic expression of ANN5 also induced the accumulation of translationally repressed RNA in plant somatic tissues. Mechanistically, ANN5 interacts with histone deacetylase 2B (HD2B) and nucleolin (NUC1), which are known epigenetic modulators of ribosome biogenesis. In metabolically active cells, ANN5 associates with ribosomal and ribosome-associated proteins, suggesting it modulates the regulatory landscape of translation. We propose that the primary physiological role of ANN5 is to coordinate the formation and subsequent availability of RNA reserves during pollen maturation and activation.

## Introduction

Mature pollen grains function independently. They achieve physiological autonomy from the maternal plant during late developmental stages, entering a state of physiological dormancy ^1^. Following hydration on the stigma, pollen grains become metabolically active to initiate germination. Rapid pollen tube emergency is strictly contingent upon the immediate initiation of translation and protein synthesis. This energy-intensive process draws on energy-storage reservoirs, such as lipids and soluble carbohydrates, that are activated during pollen hydration and subsequently catabolized^2^. Thus, during pollen maturation, large amounts of diverse RNA species accumulate alongside these essential energy reserves. The study by Hafidh et al. (2018) introduced and characterized the pollen sequestrome in *Nicotiana tabacum*, identifying it as a distinct subcellular compartment that stores and translationally represses specific mRNA transcripts in mature pollen and pollen tubes. This compartment behaves dynamically and remains distinct from actively translating polysomes and other RNA-containing fractions. The pollen sequestrome consists of non-translating single ribosomes (monosomes) that mediate mRNA sequestration during pollen maturation, activation, and pollen tube growth. Consistently, pollen germination largely depends on stored mRNAs and active protein biosynthesis rather than de novo transcription. Indeed, pharmacological studies on *Pinus bungeana* pollen grains revealed that germination is unaffected by actinomycin D (an inhibitor of mRNA transcription) while it is inhibited by cycloheximide (an inhibitor of protein synthesis) ^3^.

Pollen maturation requires strict regulation of ribosome biogenesis to establish an RNA sequestrome. In Arabidopsis, ribosomal RNA (rRNA) genes are organized as tandem repeats within nucleolar organizer regions (NORs) on chromosomes 2 and 4 ^4^. While somatic tissues actively transcribe only specific rDNA variants (VAR2 and VAR3) and silence others (VAR1 and VAR4), pollen actively transcribes all primary variants ^5,6^. Nucleolin, primarily NUC1, plays a pivotal role in this process by modulating chromatin accessibility and promoting pre-rRNA cleavage ^5,7–9^. Conversely, histone deacetylases act antagonistically to compact chromatin and repress rDNA transcription. However, like the HD2B/C heterocomplex, they can also function non-canonically in pre-rRNA processing by binding rRNA and snoRNAs to regulate methylation. Although some plant histone deacetylases lack confirmed in vitro enzymatic activity, complexes like HD2B/RPS6B efficiently repress rDNA transcription in vivo ^10–12^.

Annexins are ubiquitous proteins, typically recognized for their involvement in membrane repair and trafficking, including vesicle organization ^13–17^. The ability of annexins to bind phospholipids in a calcium-dependent manner gains new relevance in the context of membrane contact site formation during inter-organelle communication ^18^. By forming inter-organelle connections, annexins are shaping the intracellular communication network. Annexin-dependent tethering is not limited to membrane-bound structures. Binding of ANXA11 to RNA granules, and the subsequent association of these granules with lysosomes to facilitate their movement within the zebrafish axons—indicates that annexins play a role in spatial coordination of intracellular processes ^19^. In recent years, it has become evident that RNA-binding, particularly mRNA binding, is another common feature of many annexins ^20,21^. A detailed in vitro study on ANXA2 revealed that it post-transcriptionally regulates c-myc expression via interaction with both the 5 I and 3 I UTRs ^22–24^.

Previously, we showed a significant decline in the size, viability, and vigor of mature pollen in ANN5 knockdowns, resulting in an initial delay in pollen tube growth within the pistil ^25^. Our current findings implicate ANN5 in the organization of RNA reserves in maturing pollen. Here we show that ANN5 is a nucleolar protein that affects ribosomal RNA biogenesis and modulates its turnover. This in combination with our previous study unravels the multifaceted role of ANN5 in pollen biology.

## Results

### ANN5 preferentially targets the nucleolus of maturing pollen

To characterize the spatiotemporal distribution of ANN5 during late pollen development, we monitored its localization from bicellular grains to pollen maturity. Confocal imaging was performed on pollen grains from Arabidopsis plants stably expressing YFP-tagged ANN5 under the pollen-specific LAT52 promoter ^26^. The LAT52 promoter mirrors the temporal profile of the native ANN5 promoter while driving robust expression necessary for high-resolution imaging.^27^.

In bicellular pollen, ANN5-YFP predominantly localized to the nucleolus, with faint signals in the nucleoplasm and cytoplasm of the vegetative cell (Fig.1a). As pollen matured, fluorescence intensity increased in both the nucleoplasm and cytoplasm. The cytoplasmic distribution of ANN5 transitioned from distinct punctate foci in bicellular and tricellular grains into a homogeneous, continuous network. Meanwhile, the initially pronounced nucleolar distribution faded, mirroring the metabolic repression and structural condensation of the nucleolus in mature pollen.

**Figure 1.**
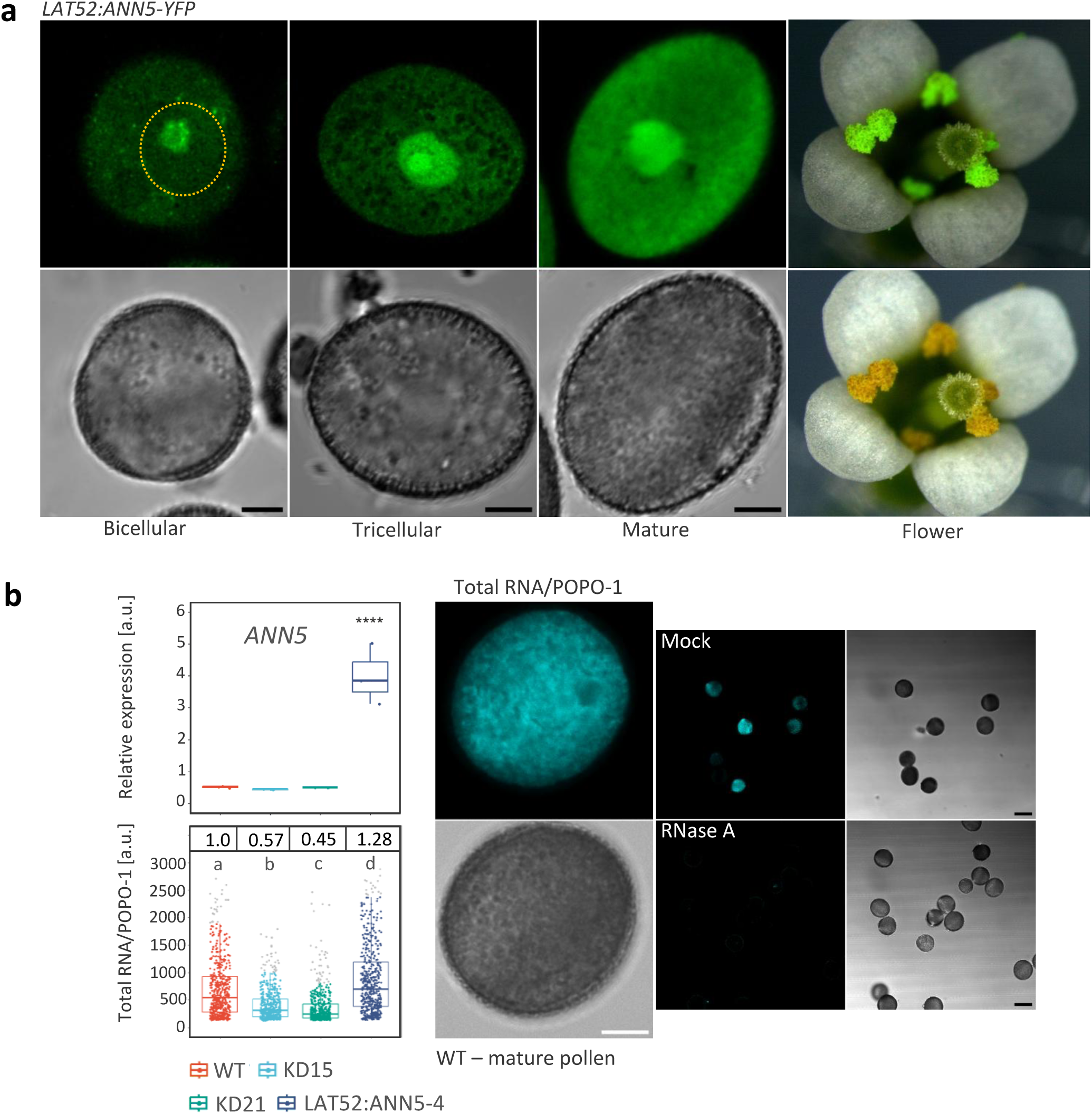
Nucleolar dominance of ANN5 during Arabidopsis pollen maturation. **a.** Representative confocal images of individual pollen grains isolated from buds of LAT52: ANN5-YFP plants capturing the nucleolar plane view. The yellow dashed oval depicts the position of the vegetative nucleus. On the right, projection along the optical axis Z of an open flower imaged with a fluorescent stereomicroscope. b. Size of pollen RNA reserves correlates with ANN5 transcript abundance. An upper box plot illustrates the ANN5 relative expression level in hydrated pollen normalized to *UBC21, RCE1, UBC9*. Differences compared with WT were assessed by ANOVA, followed by Tukey’s HSD test. Only significant differences are marked; **** indicates p-value < 0.0001. For each group, n = 3. The lower box plot illustrates the POPO-1 RNA-specific dye relative fluorescence intensity in individual pollen grains from wild-type, *ANN5* knockdowns (KD15, KD21), and the *LAT52: ANN5-YFP* line 4. Different letters (a, b, c, d) indicate statistically significant differences between groups as determined by Kruskal-Wallis followed by Dunn’s post hoc test. Outliers are marked in grey. For each group, n = 500. Representative image “Total RNA/POPO-1“ image illustrates the compartmentalization of pollen RNA reserves. Scale bar = 5 µm. The “Mock/RNaseA” images demonstrate the RNA specificity of the POPO-1 dye, which is completely eliminated following a 1-hour RNaseA treatment. Confocal images were acquired at the same settings. Scale bar = 20 µm.

Mature pollen grains contain large amounts of RNA, mostly rRNA. We assessed whether ANN5 affects rRNA accumulation by quantifying nucleic acid dye incorporation in single pollen grains using POPO-1 iodide (Extended Data Fig.1). Staining revealed that pollen RNA reserves were compartmentalized into a distinct branched network rather than homogeneously dispersed (Fig.1b). Specificity was confirmed via RNase treatment, which abolished the signal (Fig.1b). Compared to the control, total RNA levels were significantly reduced by 40-50% in pollen from *ANN5* RNAi-silenced lines KD15 and KD21. Conversely, ANN5 overexpression (LAT52:ANN5-YFP line 4) increased RNA abundance by over 20%. These findings suggest that ANN5 regulates RNA accumulation, particularly rRNA, during pollen development.

**Table 1.**
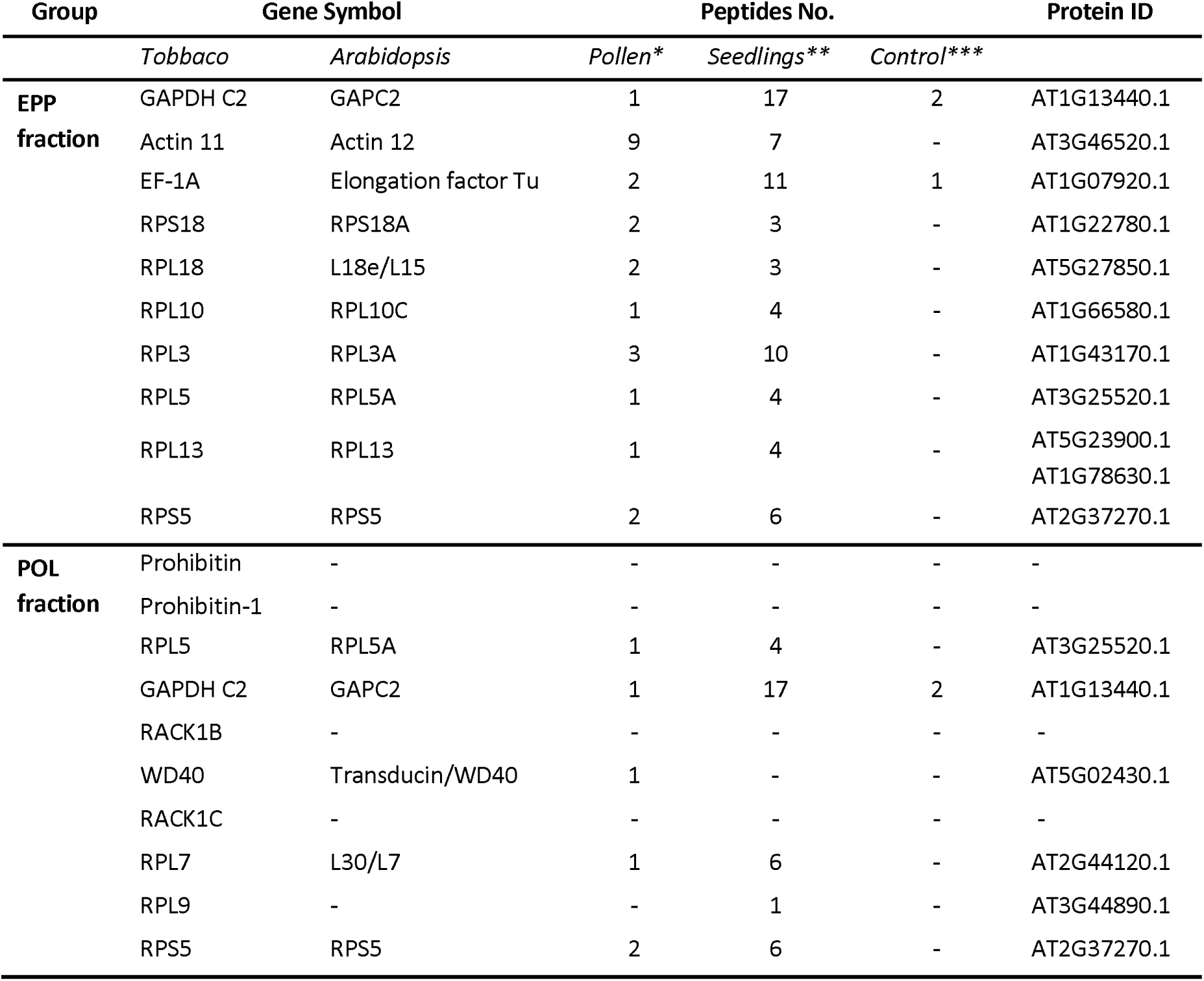
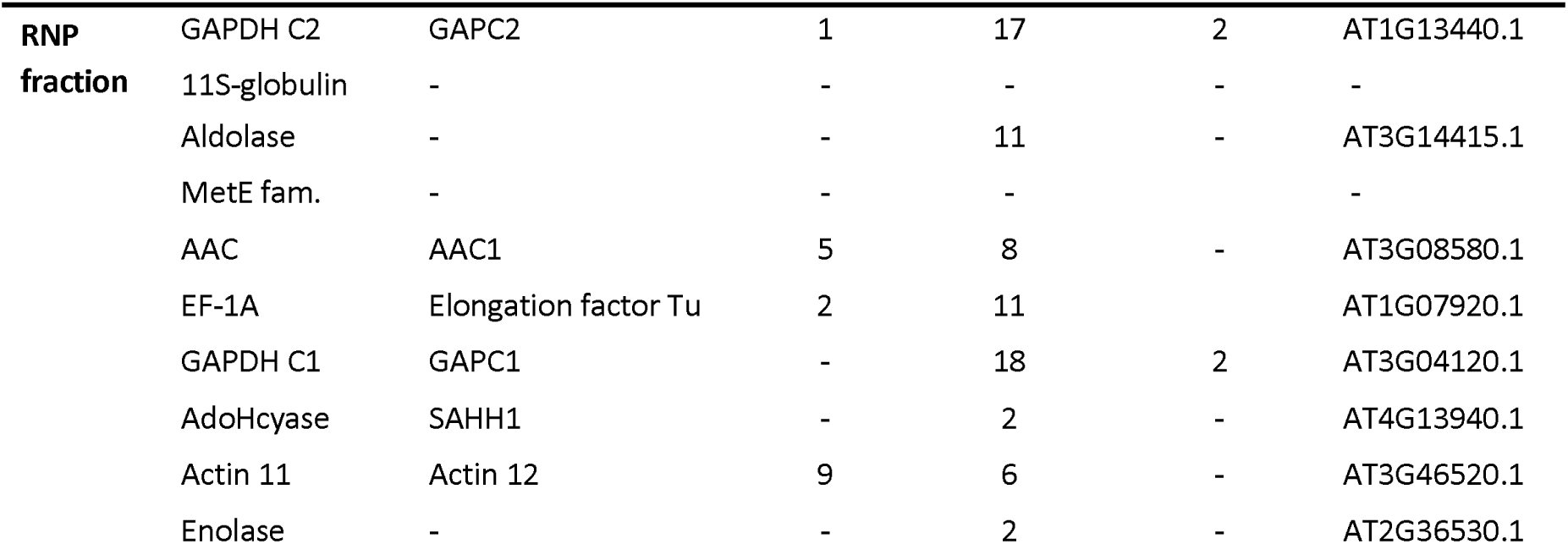
ANN5 co-purifies with the translational machinery of the pollen sequestrome. Proteins co-purified with ANN5 were identified by mass spectrometry. They were categorized into three functional groups (EPP, POL, and RNP fraction) based on the tobacco pollen RNA fraction classification by Hafidh et al. (2018). The EPP fraction corresponds to EDTA/ puromycin-resistant particles for long-term storage of transcripts, translating polysomes constitute the POL fraction, and free ribonucleoprotein particles, the RNP fraction. ANN5 was pulled down from hydrated pollen of Arabidopsis LAT52:ANN5-YFP_4 (*) and 7-day-old seedlings overexpressing ANN5, line OE2 (**). Arabidopsis seedlings expressing 35S:GFP were used as a control sample (***). Proteomic data are provided as Supplementary Data 1.

### Ectopic expression of ANN5 triggers the accumulation of a translationally repressed RNA in seedlings

To study the impact of ANN5 on early ribosome biogenesis stages, we utilized 7-day-old Arabidopsis seedlings stably overexpressing ANN5. In the OE2 line, ANN5 expression is driven by a double 35S promoter, whereas the OE3 line utilizes a single 35S promoter, resulting in proportionally elevated ANN5 transcript levels and larger seedlings (Fig.2a). Confocal microscopy revealed that ANN5-YFP accumulated predominantly in the nucleoli of primary root cells in both OE lines (Extended Data Fig.2), validating them for further investigation. While the level of endogenous ANN5 transcripts in wild-type seedlings was exceptionally low, alterations in ANN5 expression resulted in a distinct phenotype characterized by reduced/increased seedling size (Fig.2a), consistent with previous reports ^25^.

**Figure 2.**
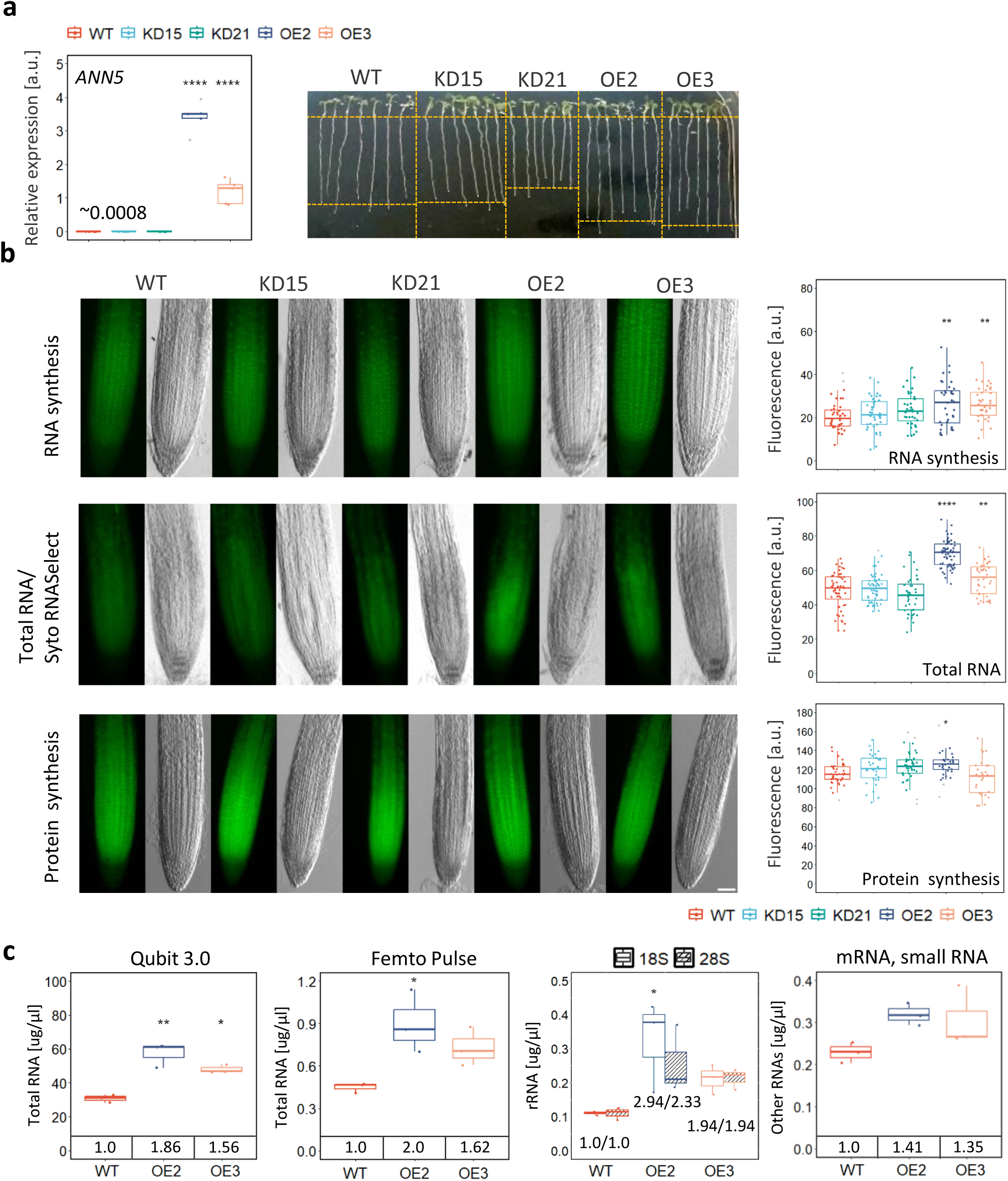
Ectopic expression of ANN5 drives translationally inactive RNA accumulation in somatic tissues. **a.** Representative image of 7-day-old seedlings of wild-type (WT), ANN5 knockdowns (KD15, KD21), and ANN5 overexpressing lines: OE2 and OE3 growing on MS-agar plates with 1% sucrose. The box plot shows ANN5 relative expression in 7-day-old seedlings, normalized to YLS8 and UBC21. For each genotype, n = 5; outliers are marked in grey. b. Images of individual root tips (green channel and brightfield) were captured with a fluorescent stereomicroscope. Incorporation of 5-ethynyl uridine (EU) or L-azidohomoalanine (AHA) was detected with Alexa Fluor 488 azide or an alkyne variant of the dye. Total RNA was stained with Syto RNASelect dye. Scale bar = 20 µm. Box plots on the right illustrate relative fluorescence intensity measured in the root tips. n = 25-63. **c.** RNA quantification and identification by pulse field capillary electrophoresis. Box plots show RNA amount and content (µg/µl) isolated from 20 wild-type seedlings and the ANN5-OE2 line (measured using Qubit 3.0) and after dilution (measured using Femto Pulse). The values displayed inside the plot frame at the bottom indicate relative changes in RNA accumulation in the transgenic lines compared to WT. Source data are provided as Extended Data Fig.4. For all assays, differences relative to WT were evaluated by ANOVA, followed by Tukey’s post hoc HSD test. Significance is indicated as follows: p < 0.05, ** p < 0.01, and **** p < 0.0001.

First, we assessed de novo RNA synthesis rates by incubating seedlings (WT, KD15, KD21, OE2, and OE3) in liquid MS medium with 5-ethynyl uridine (EU) for 90 minutes. Nascent transcripts were detected via click chemistry with a modified Alexa Fluor 488 dye. Fluorescence stereomicroscopy showed extensive EU incorporation in root apex cells (Fig.2b). Because most newly synthesized RNA is ribosomal, the signal localized predominantly within the nucleolus ^28^. Measurements revealed approximately 25% higher fluorescence intensity in the root apices of ANN5-OE lines than in controls, indicating that ANN5 promotes de novo rRNA synthesis.

To quantify steady-state total RNA levels, methanol-fixed seedlings were stained with Syto RNASelect. Total RNA levels in root apices were significantly higher in ANN5-OE lines, showing a 40% increase in the OE2 line compared to controls (Fig.2b). RNase treatment confirmed staining specificity (Extended Data Fig.3). Next, total RNA from WT and ANN5-OE seedlings was separated by pulse-field capillary electrophoresis. This analysis revealed a significant increase in 18S and 28S rRNA in ANN5-OE lines (Fig.2c, Extended Data Fig.4), demonstrating that ANN5 modulates rRNA abundance in a dose-dependent manner.

We next investigated whether excess rRNA influences overall protein synthesis using a click chemistry assay with the methionine analog L-azidohomoalanine (AHA). The abundance of newly formed peptides remained nearly equal across all genotypes, except for a slight increase in the OE2 line. This suggests that not all accumulated rRNA in ANN5-OE lines is translationally active (Fig.2b). Cycloheximide pre-incubation confirmed AHA specificity (Extended Data Fig.3).

### ANN5 associates with the free ribosomal subunits and a subpopulation of monosomes

To determine the specific stage of ribosome biogenesis involving ANN5, we employed GFP-TRAP affinity purification followed by LC-MS/MS proteomics on Arabidopsis seedlings (35S:ANN5-OE2) and hydrated mature pollen (LAT52:ANN5-YFP). This revealed essential early-stage rRNA biogenesis factors, including histone deacetylase 2B (HD2B) and nucleolin (NUC1), in ANN5-overexpressing seedlings. These interactors were absent in hydrated mature pollen, where ribosome biogenesis is complete. A large subset of ribosomal proteins co-purified with ANN5 from both seedlings and hydrated pollen. We screened our proteomic data for Arabidopsis homologs of tobacco pollen proteins assigned to three distinct RNP populations ^29^. Notably, all homologs characteristic of the translationally repressed RNA fraction (pollen sequestrome) co-purified with ANN5 from both tissues, with higher peptide abundance in seedlings. These proteins did not co-purify with the free GFP control (Tab.1).

To determine the physical distribution of ANN5 across ribosomal fractions, cycloheximide and chloramphenicol treated extracts from seedlings and mature pollen were separated on a sucrose density gradient. Western blot analysis showed that ANN5 was most abundant in fractions containing free 40S and 60S ribosomal subunits (Fig.3a). In hydrated pollen, single ribosomes (monosomes) were prevalent, while heavy polysomes were absent. Intriguingly, ANN5 associated with a specific subpool of monosomes, rather than the entire monosomal pool. A similar distribution pattern of ANN5 across ribosomal species was observed in seedlings, with heavy polysomes largely devoid of ANN5. The polysome profiles for wild-type and OE2 seedlings were virtually identical, corroborating our in planta protein synthesis assay (Fig.2b) and indicating comparable global translational activity (Extended Data Fig.5).

**Figure 3.**
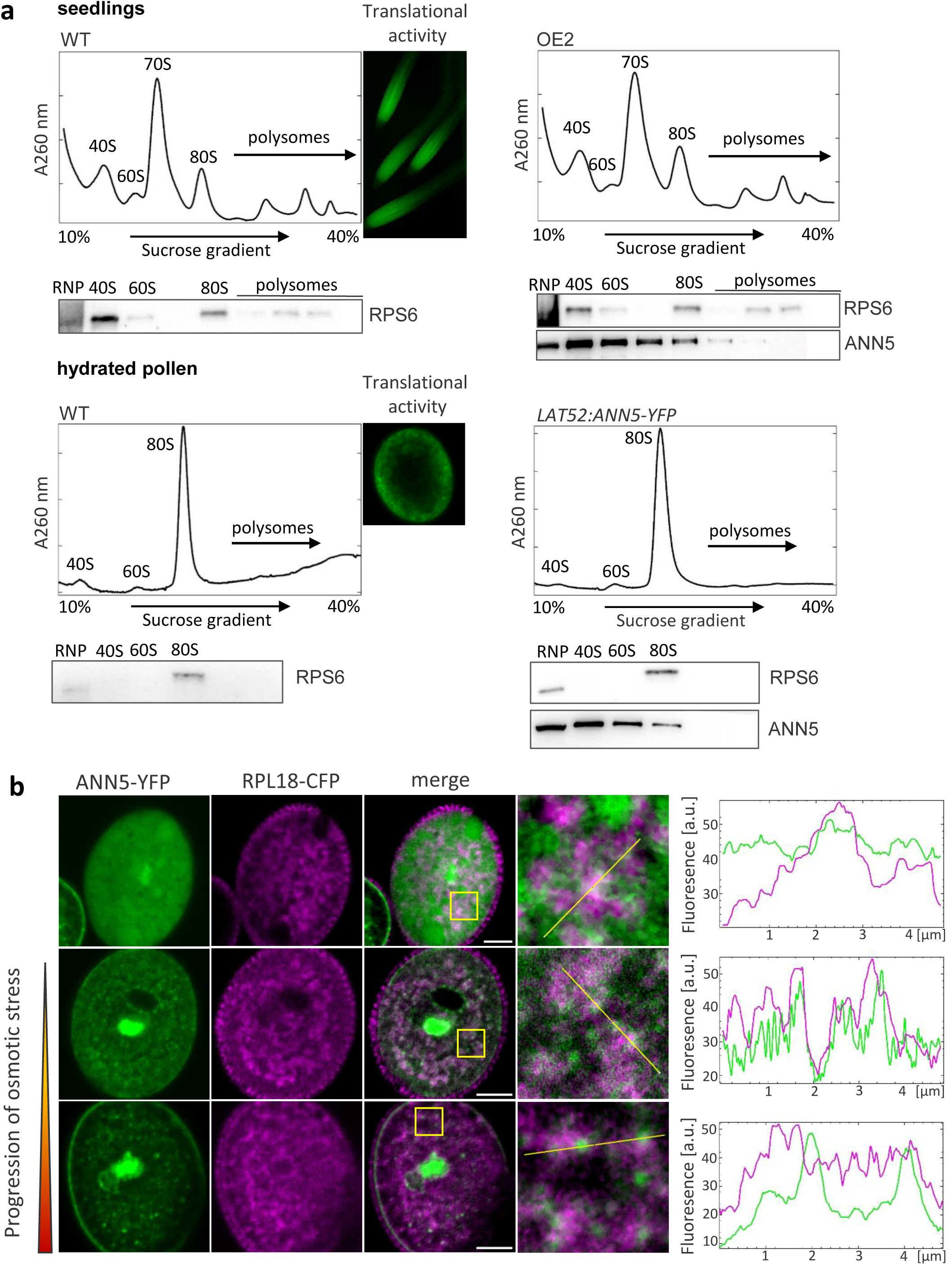
ANN5 co-migrates with free 40S and 60S ribosomal subunits and 80S monosomes on a sucrose density gradient. **a.** Polysome profiling by sucrose density gradient centrifugation of 7-day-old seedlings (OE2) and hydrated pollen of wild-type and overexpressing ANN5. Eluted fractions were run on Western blots and probed with PRS6 and GFP antibodies. Representative images illustrate translational activity in seedlings and hydrated pollen after 1,5/1 h of incubation with modified L-azidohomoalanine (AHA), detected using the Alexa Fluor 488 alkyne dye. **b.** Single confocal images show subcellular distribution of ANN5-YFP and RPL18-CFP in hydrated pollen. Images were acquired 10 min (upper panel), 20 min (middle panel), and 30 min (lower panel) after pollen hydration. Graphs illustrate different scenarios for ANN5 distribution in the ribosome-rich area. Scale bar = 5 µm.

To explore the spatial interplay between ANN5 and ribosomes in mature pollen, we generated Arabidopsis lines expressing ANN5-YFP and RPL18-CFP, using the latter as a marker for the large 60S subunit. In hydrated mature pollen, ribosomes tagged with RPL18-CFP occupied a large volume of the vegetative cell cytoplasm, forming an extensive network of clusters. Defining the precise association between ANN5 and ribosomes was challenging in most pollen grains because ANN5 exhibited a distribution pattern largely overlapping with RPL18 (Fig.3b). We previously reported that ANN5 relocated from the cytosol to the pollen membrane (PM) during prolonged hydration, leading to PM rupture ^26^. In such grains, a pool of ANN5 remained attached to intracellular structures in the central vegetative cell. Here, we showed that these cytoplasmic structures contain ribosomes. Moreover, ANN5 formed distinct assemblies attached to the boundaries of RPL18-tagged ribosome clusters, which grew as stress persisted.

### Both the transcription and processing of pre-rRNA are influenced by ANN5

NUC1 and HD2B, which co-purified with ANN5 from seedlings, act at the earliest stages of rRNA biogenesis, participating in both rDNA transcription and the initial steps of 45S pre-rRNA processing. HD2B forms a complex with RPS6B in the nucleolus to repress rDNA transcription ^30^. Since ANN5 co-purified with RPS6, which has two highly homologous paralogs (RPS6A and RPS6B), we tested both variants for ANN5 interactions. We combined BiFC (bimolecular fluorescence complementation) and GFP-TRAP co-purification to verify protein-protein interactions in planta. Concurrently, RT-PCR profiled rRNA allele expression, while Northern blotting assessed 45S pre-rRNA processing kinetics.

Gene constructs encoding fluorescently tagged proteins were transiently expressed in N. benthamiana leaves. All tested proteins (HD2B, RPS6A, RPS6B, NUC1) localized preferentially to the nucleolus and nucleoplasm, though ANN5 and the RPS6 paralogs also showed a distinct cytoplasmic presence (Fig.4a). Nucleolar redistribution of ANN5 was most pronounced upon co-expression with RPS6A and NUC1. A complementary BiFC assay revealed that ANN5 interacts with HD2B and NUC1 exclusively within the nucleolus (Fig.4a). Unlike HD2B, which associates with both RPS6 paralogs in the nucleolus, ANN5 formed BiFC complexes exclusively with RPS6B. Notably, ANN5-RPS6B BiFC complexes localized to both the nucleolus and cytoplasm, whereas no signal was detected for RPS6A (Fig.4a). GFP-TRAP assays confirmed that ANN5 interacts predominantly with HD2B and exhibits a strict binding preference for RPS6B over RPS6A (Fig.4b).

**Figure 4.**
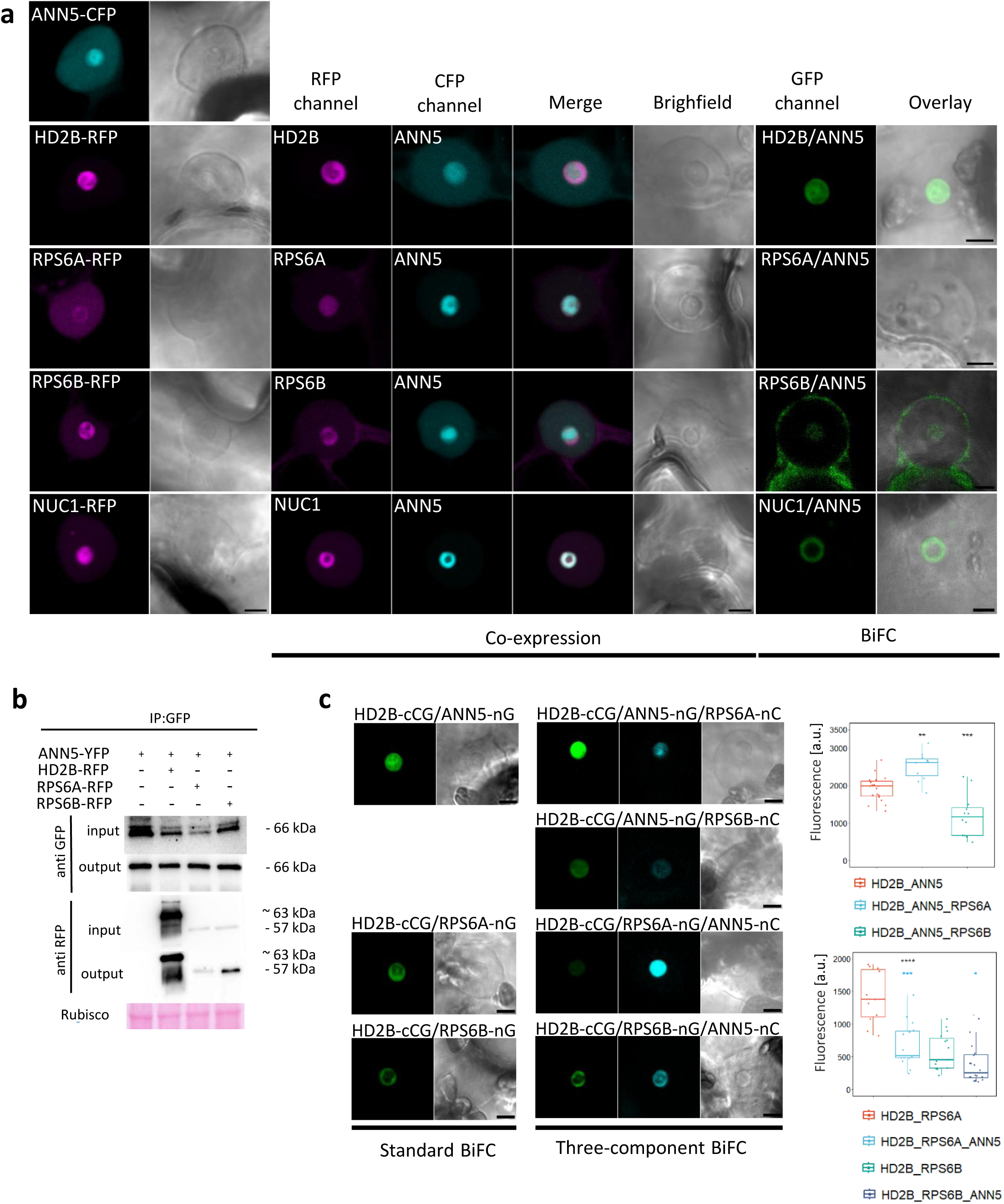
ANN5 interacts with HD2B and NUC1, and modulates the organization of the HD2B/RPS6 complex in the nucleolus. **a.** *N. benthamiana* epidermal cells transiently expressing ANN5 (cyan) or/and HD2B, RPS6A, RPS6B, and NUC1 (magenta). Single confocal images of the nucleus/nucleolus plane illustrate the combination of co-expression and GFP BiFC assays. On the left are control images showing each protein expressed individually. Scale bar = 10 µm. **b.** GFP-Trap co-immunoprecipitation analysis reveals that ANN5-YFP preferentially interacts with HD2B-RFP and RPS6B-RFP over RPS6A-RFP. Blots were probed using anti-GFP and anti-RFP antibodies. Ponceau staining served as a loading control. Representative results from three independent biological replicates are shown. c. Confocal images of the nucleus/nucleolus plane illustrate results from the comparative study using standard and three-component BiFC assays. The upper box plot illustrates changes in HD2B/ANN5 BiFC complex formation induced by nC-tagged RPS6A or RPS6B proteins, as monitored in the GFP channel. n = 11-19. Differences between three-component BiFC with HD2B/ANN5 were assessed using ANOVA, followed by Tukey’s HSD test. Only significant differences are marked; ** indicates p-value < 0.01, *** indicates p-value < 0.001. The lower box plot illustrates how the nC-tagged ANN5 alters the formation of HD2B/RPS6A or HD2B/RPS6B BiFC complexes, as monitored in the GFP channel. n = 11-20. Differences between the triple-component combination and the double-component combination were assessed using the Kruskal-Wallis test, followed by the Wilcoxon test. *** indicate p-value < 0.001, * indicate p-value < 0.05.

Next, a three-component BiFC assay evaluated competitive interactions between a shared bait and two potential interactors ^31^, enabling us to examine the interplay among HD2B, RPS6, and ANN5. Upon co-expression, GFP and CFP were reconstituted simultaneously. First, we monitored GFP reconstitution for HD2B/RPS6 complexes. The presence of ANN5 decreased GFP signal intensity regardless of the RPS6 paralog used, indicating it affects HD2B/RPS6 complex formation (Fig.4c). Second, we assessed HD2B/ANN5 complex formation with or without RPS6 paralogs (Fig.4c). RPS6A increased GFP signal intensity beyond the HD2B/ANN5 control, suggesting that ANN5 and RPS6A cooperatively bind HD2B, with RPS6A acting as an allosteric modulator. Conversely, introducing RPS6B decreased the HD2B/ANN5 BiFC signal, likely because ANN5 traps a pool of RPS6B in the cytoplasm, reducing their nucleolar availability for HD2B. Thus, ANN5 alters the organization of this multimeric complex when associated with HD2B, and its cytoplasmic interaction with RPS6B suggests it may affect non-nucleolar roles of RPS6B.

We then examined the rRNA allele expression profile in seedlings with modified ANN5 levels. Using RT-PCR with P3 and P4 primers targeting the 3’ETS region, we examined 45S pre-rRNA transcription^5^. Length variations in these sequences enable the identification of four rRNA variants (VAR1–4). All four variants were detected, with VAR2 and VAR3 being most prevalent. However, VAR2 pre-45S rRNA levels decreased in ANN5-OE lines compared to wild-type seedlings (Fig.5a). Notably, an additional rRNA subvariant was upregulated in the OE2 line, suggesting that ANN5 may orchestrate ribosome heterogeneity by altering individual 45S pre-rRNA variants.

**Figure 5.**
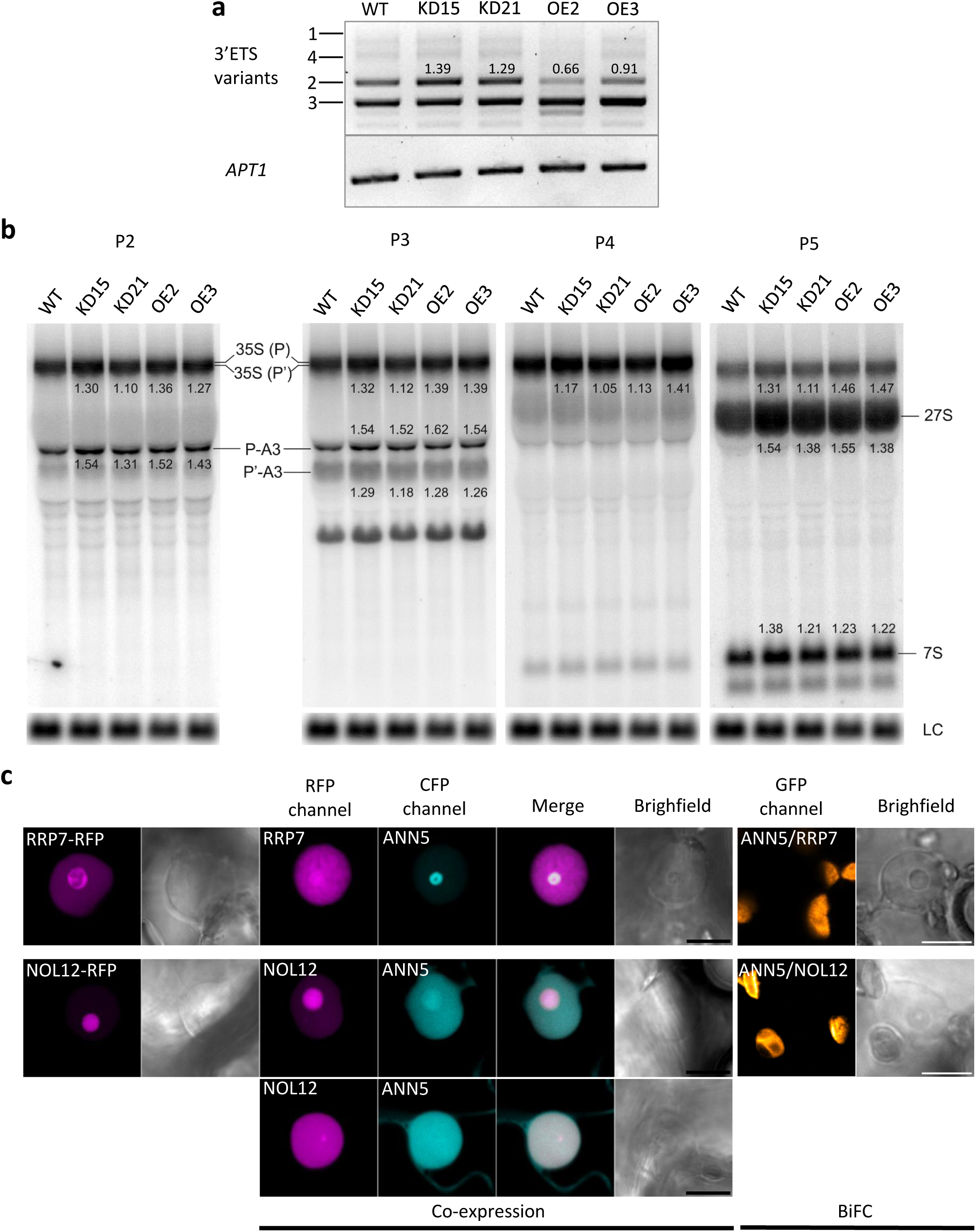
Alterations in the expression of 3’ETS rDNA variants, pre-rRNA processing efficiency, and the subcellular distribution of pre-rRNA processing factors RRP7 and NOL12 all correlate with ANN5 expression levels. **a.** RT-PCR with p3, p4 primers and cDNA prepared from WT, KD15, KD21, OE2, and OE3 plants. The amount of cDNA was assessed with APT1 primers in a control reaction. The values show changes in VAR2 transcript accumulation in the transgenic lines compared to WT. **b.** Northern blot analysis of rRNA precursors and intermediates. RNA isolated from 7-day-old Arabidopsis seedlings was separated on 1.1% agarose and hybridized with probes P2, P3, P4, and P5. rRNA precursors and intermediates are described on the right. The values shown above or below the RNA intermediates indicate changes in their accumulation in the transgenic lines compared to WT. U2 snRNA was used as a loading control (LC). The analyses were performed in two biological replicates. c. Representative single confocal images of the nucleus/nucleolus plane illustrate the results of ANN5 co-expression with RRP7-RFP or NOL12-RFP and BiFC assay in *N. benthamiana* epidermal cells. For ANN5 and NOL12, two representative scenarios illustrating their mutual influence are presented. Scale bar = 10.

**Figure 6.**
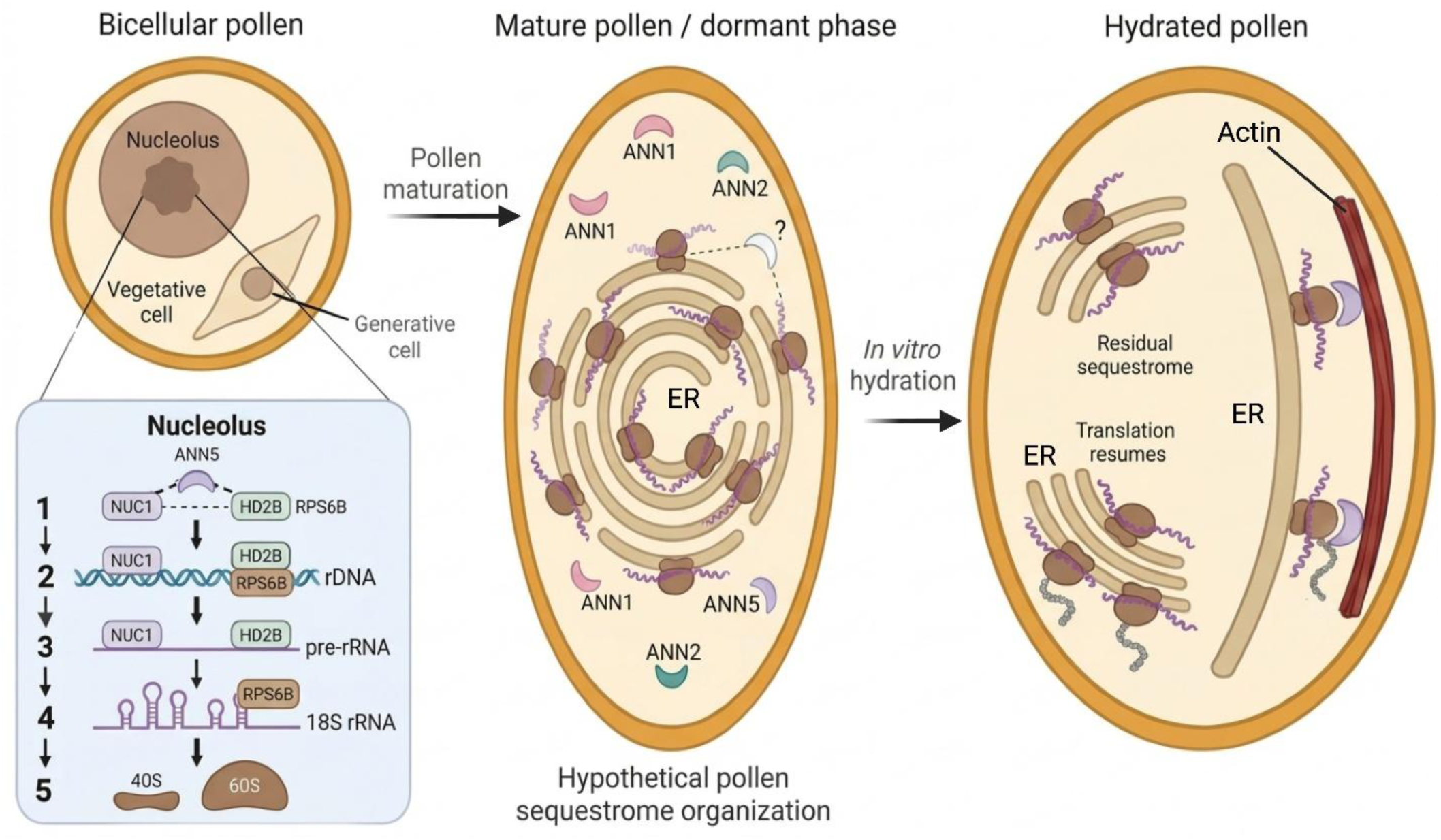
Schematic diagram illustrating the dual role of ANN5 in rRNA metabolism in Arabidopsis pollen. In this model, ANN5 acts first as an architect of the translationally inactive sequestrome during maturation, and subsequently transitions into a dynamic modulator of the ribosomal environment upon hydration. **Pollen maturation**: At the bicellular stage, ANN5 modulates nucleolar dynamics by associating with nucleolin 1 (NUC1) and histone deacetylase 2B (HD2B) to regulate rDNA transcription and pre-rRNA processing. This ANN5 activity is required for the accumulation of RNA reserves within the pollen sequestrome. Upon reaching maturity, the pollen grain enters dormancy. Based on available data, we propose a hypothetical model for the organization of this sequestrome in mature pollen ^29,49,60^. However, the specific annexins that associate with the pollen sequestrome during developmentally induced dormancy remain to be identified. *Pollen activation*: Upon hydration, translational activity resumes, and ANN5 acts as a molecular anchor that tethers a specific subset of monosomes to the actin cytoskeleton. This molecular linkage facilitates the intracellular redistribution of distinct components of the pollen sequestrome within the activated vegetative cell. The precise identity of the mRNAs loaded onto these ANN5-associated ribosomes remains to be elucidated.

To investigate ANN5 involvement in 45S pre-rRNA processing, we analyzed rRNA precursor patterns and levels via Northern blotting total RNA from 7-day-old seedlings (WT, KD15, KD21, OE2, OE3). Both knockdown and overexpression lines exhibited mild alterations in 45S pre-rRNA processing, with KD15 showing the most pronounced accumulation of unprocessed intermediates (Fig.5b). These results clearly illustrate that pre-rRNA processing is sensitive to the precise dosage of ANN5.

NUC1 and HD2B facilitate the recruitment of snoRNAs required for early pre-rRNA processing ^7,10^. To visualize the processing alterations shown by Northern blots, we co-expressed ANN5 with RRP7 or NOL12, processome factors acting downstream of NUC1 and HD2B, in *N. benthamiana* leaves ^32–34^. When expressed individually, RRP7 and NOL12 localized within a specific nucleolar subregion and the nucleoplasm. Co-expression with ANN5 disrupted this, shifting both proteins toward the nucleoplasm due to severely altered nucleolar organization. This shift indicates enhanced nuclear retention of processing factors, possibly reflecting localized rRNA deposition. Crucially, neither RPL17 nor NOL12 co-purified or interacted with ANN5 in BiFC assays (Fig.5c). Combined, our findings demonstrate that ANN5 accelerates rDNA transcription, causing a subset of rRNA to become trapped in the nucleus during pre-rRNA processing.

## Discussion

Upon maturity, pollen grains enter developmental arrest and cease active biosynthesis, a process preceded by the accumulation of non-translating monosomes and translationally repressed mRNAs within the pollen sequestrome ^1,29^. While similar mRNA storage strategies occur during seed desiccation ^35^, the molecular mechanisms driving pollen RNA sequestration have remained elusive. We demonstrate that ANN5 triggers the accumulation of translationally inactive ribosomal RNA (rRNA), a core constituent of the pollen sequestrome. This pathway operates natively during Arabidopsis pollen maturation and can be ectopically recapitulated in vegetative tissues, indicating that ANN5 is sufficient to reprogram somatic translational machinery from active utilization to RNA storage.

Nucleolar association of ANN5 with HD2B and NUC1 suggests a multi-level regulation of ribosome biogenesis, encompassing rDNA transcription and pre-rRNA processing factor recruitment ^7,10^. Given that annexins modulate interaction partners via steric hindrance, co-receptor scaffolding, or induced proximity ^36–39^, and can link to chromatin-remodeling complexes ^40–42^, we propose that ANN5 allosterically modulates HD2B and NUC1 to alter rDNA transcription at the onset of pollen maturation.

Ectopic *ANN5* expression in somatic tissues markedly increases steady-state levels of translationally inactive rRNA. While the majority of this surplus RNA undergoes correct processing, intensive ANN5-driven synthesis triggers nucleolar and proteostatic stress ^43,44^. This overproduction overwhelms endogenous processing pathways, as evidenced by our Northern blot analysis showing accumulation of pre-rRNA intermediates in *ANN5*-OE lines. These excess intermediates likely obstruct nuclear export, promoting nucleolar and nuclear RNA retention ^45^, a model supported by the nucleolar reorganization and nucleoplasmic relocation of RRP7 and NOL12 observed in *N. benthamiana*. Despite these alterations, click chemistry protein synthesis assays and polysome profiling revealed unaltered translational activity in *ANN5*-OE lines, confirming that homeostatic cytoplasmic RNA levels are maintained while excess RNA is sequestered within the nucleolus and nucleus ^46–48^. Notably, ANN5 also functions as a bona fide ribosome-associated protein that specifically interacts with a translationally repressed ribosomal subpopulation, outlining a dual role for ANN5 in both RNA sequestration and ribosome-specific translational arrest.

While Arabidopsis mature pollen expresses three annexins: ANN1, ANN2, and ANN5 ^49^, their structural association with RNA-containing fractions upon pollen dormancy has been unclear. In tobacco, the ANN5 homolog is absent from RNA fractions in dormant pollen but associates weakly with the pollen sequestrome and polysomes upon hydration and germination, whereas the tobacco ANN2 homolog shifts from the pollen sequestrome to free ribonucleoprotein complexes (RNPs) during this transition. Consistently, we found that Arabidopsis ANN5 co-purifies with ribosomal and ribosome-associated proteins of the pollen sequestrome upon pollen hydration. Analogous to the tobacco homolog, ANN5 may bypass RNA fractions in dormant pollen and instead modulate the ribosomal regulatory environment upon hydration. This is supported by the dynamic condensation of ANN5 in the immediate vicinity of ribosomes, visualized via the RPL18-CFP marker under prolonged osmotic stress, which may form a protective physical hub shielding cellular components during stress-induced translational arrest ^50^.

The association of tobacco RNA reserves (sequestrome and RNPs) with Actin 11, together with our finding that ANN5 specifically interacts with Actin 12 and the pollen sequestrome, suggests a mechanism in which specialized factors bridge monosomes and RNPs to distinct actin structures during pollen dormancy and hydration. Given that specific annexins bind and bundle actin to anchor mRNA subpopulations to the cytoskeleton ^51–54^, this coupling of ribosomes with distinct actin isoforms likely directs the spatial assembly of translational hubs required for pollen germination and tube elongation. Similar to how mammalian ANXA11 anchors RNA granules to moving lysosomes ^19^, ANN5 may tether ribosomes to intracellular transport machineries to facilitate their relocation and long-distance transport during pollen hydration and pollen tube emergence.

In conclusion, we propose a model wherein ANN5 orchestrates pollen sequestrome assembly and defines the ribosomal regulatory landscape. During the bicellular stage, ANN5 drives ribosome heterogeneity by interacting with NUC1 and HD2B to modulate rRNA synthesis and allele expression profiles. Its preference for specific ribosomal protein paralogs indicates selective association with a distinct ribosome subpopulation during pollen maturation or hydration. Upon hydration, ANN5 coordinates its actin-binding capacity to anchor silenced or pre-activated monosomes at subcellular sites designated for the translation of stored mRNAs. Deciphering the exact identity of these ANN5-associated mRNAs and the precise regulatory impact on their translational efficiency represents an essential next step.

## Methods

### Plant material and growth conditions

The experiments were carried out on *Arabidopsis thaliana* Col-0 accession and *Nicotiana benthamiana* plants. Arabidopsis lines with modified ANN5 levels were described previously or characterized in this study ^25,26^. We used two ANN5 RNAi-silenced lines: KD15 and additional line KD21, T5 and T3 generation, respectively. We used three lines overexpressing ANN5 under the control of the 35S promoter: OE2, OE3, and the pollen-specific promoter LAT52 (line 4). ANN5-OE3 line was generated in this study by floral-dipping transformation of the Arabidopsis ^55^ For Arabidopsis transformation, the full-length ANN5 cDNA cloned into the pGWB441 binary vector (fused to YFP) was introduced into *Agrobacterium tumefaciens* strain GV3101 (pMP90). Generation of lines overexpressing ANN5 used in this study: T5 – OE2, T3 – OE3, T3 - *pLAT52:ANN5-YFP* line 4. Arabidopsis seedling cultures were maintained on the solidified Murashige and Skoog (MS) medium (Duchefa, Amsterdam, The Netherlands) containing 1% (w/v) sucrose and 1% (w/v) agar (Duchefa, Amsterdam, The Netherlands) in controlled-environment chambers at 22 °C under 8 h of light. Arabidopsis plants were grown in Jiffy7 pots in controlled-environment chambers (Percival Scientific, Iowa, USA) at 22°°C, ca. 55% relative humidity, under 8 h of light for 4 weeks, followed by 16 h of light conditions. *N. benthamiana* plants were grown in soil under controlled environmental conditions (21^°^C, 16 h of light).

### Plasmid constructions for transient expression in N. benthamiana

The coding sequences of *HD2B, NUC1, RPS6A, RPS6B, RRP7, and NOL12* were PCR-amplified and cloned into pENTR1A vector or using pENTR^TM^/D-TOPO^TM^ Cloning Kit (Thermo Fisher Scientific), compatible with the Gateway system. The resulting plasmid was LR recombined into GWB441, GWB444, pB4GWcCG, pB4GWnG, pB4GWnC, and the resulting binary plasmids were transformed into *Agrobacterium* strain GV3101. Subsequently, Agrobacterium cultures carrying appropriate constructs were infiltrated into leaves of *N. benthamiana*, and after 72 h, leaf samples were analyzed using laser scanning confocal microscopy.

### Click chemistry assays in seedlings

De novo RNA and protein synthesis were visualized using the Click-iT™ RNA Alexa Fluor™ 488 Imaging Kit and the Click-iT™ AHA Alexa Fluor™ 488 Protein Synthesis HCS Assay Kit (Invitrogen), respectively. Seven-day-old seedlings were incubated in liquid MS medium containing either 50 µM 5-ethynyl uridine (EU; for RNA synthesis) or 50 µM L-azidohomoalanine (AHA; for protein synthesis) for 90 min at 22°C under daylight conditions. Following incubation, seedlings were fixed in 3.7% formaldehyde and permeabilized with 0.1% Triton X-100. Fluorescent detection was performed by incubating the samples in the respective Alexa Fluor 488 reaction cocktails for 30 min. To estimate the level of non-specific binding of the Alexa Fluor 488 dye to fixed plant tissues, control samples were prepared by omitting EU and AHA during incubation in MS medium. To eliminate non-specific Alexa Fluor 488 binding from the Click-iT™ RNA Alexa Fluor™ 488 Imaging Kit, the manufacturer’s protocol was modified by adding a 3% BSA blocking step, adapted from the protein synthesis kit protocol. A one hour pre-incubation with 50 µg/mL cycloheximide was performed to validate the specificity of AHA incorporation.

### Microscopy

Root apices of *Arabidopsis* seedlings were imaged using a Leica M205 FCA fluorescence stereo microscope. Confocal images of pollen samples and *N. benthamiana* leaves expressing CFP-, GFP-, YFP-, or RFP-tagged proteins were acquired with a Nikon C1 and a Leica Stellaris 8 Falcon microscope at the Fluorescence Microscopy Facility (IBB PAS, Poland). The Nikon C1 setup utilized a 40x Plan Fluor, 60× Plan-Apochromat oil immersion objectives (Nikon Instruments), laser lines at 408, 488, and 543 nm, and emission filters of 450/35 nm (CFP, POPO-1), 515/530 nm (GFP, YFP), and 605/75 nm (RFP). Leica Stellaris 8 Falcon imaging was performed using an HC PL APO 63×/1.20 water immersion objective (Leica Microsystems) and a white-light tunable laser. For this system, excitation and emission detection windows were configured as follows: POPO-1 (ex: 405 nm, em: 430–460 nm), CFP (ex: 440 nm, em: 460–490 nm), GFP (ex: 488 nm, em: 505–530 nm), and YFP (ex: 514 nm, em: 530– 560 nm). To minimize bleed-through between fluorescence channels, low laser power and sequential single-channel acquisition were used. Confocal images were analyzed using Fiji software.

### RNA methods

#### RNA staini

For RNA labeling, seedlings and pollen grains were stained using SYTO RNASelect Green Fluorescent Cell Stain and POPO-1 iodide (Thermo Fisher Scientific). Seven-day-old seedlings were fixed in pre-chilled methanol at −20°C for 20 min and washed twice in PBS for 5 min each. Fixed seedlings were then incubated in a 500 nM SYTO RNASelect solution in PBS for 45 min at room temperature, followed by two 5-min PBS washes before imaging. Unfortunately, SYTO RNASelect dye was poorly incorporated by both methanol- and paraformaldehyde-fixed pollen. Next, we tested dimeric cyanine nucleic acid stains and selected POPO-1 iodide, which exhibited superior selectivity for total RNA over the cytoplasm compared with other dimeric cyanine dyes. Mature pollen was collected in ice-cold 0.3 M mannitol, as previously described ^57^. Following fixation in 3.7% paraformaldehyde, pollen grains were permeabilized in 0.1% Triton X-100 in PBS for 1.5 h. Total RNA was stained using 0.5 µM POPO-1 in PBS for 1 h, followed by two 5-min washes in PBS before imaging. To validate the specificity of RNA dyes incorporation, samples were treated with 1 mg/mL RNase A in PBS supplemented with 1 mM MgCl₂ at 37°C for 30–60 min after permeabilization and before staining.

#### Total RNA extraction

total RNA was isolated from 7-day-old seedlings for pulse-field capillary electrophoresis, RT-PCR, and RT-qPCR using Syngen Plant RNA Mini Kit (Syngen) and for Northern blotting using TRI Reagent (Sigma-Aldrich) according to the manufacturer’s instructions.

#### RNA pulse-field capillary electrophoresis

prior to analysis, RNA concentration was measured on Qubit 3.0. To meet the concentration requirements for pulse-field capillary electrophoresis while preserving the relative differences in total RNA yield between the Arabidopsis lines, samples were diluted proportionally using a uniform dilution factor (Extended Data Fig.4). Measurements were taken using an Agilent Femto Pulse system, and results were analyzed with ProSize 3.0 software.

#### RT-qPCR

Isolated RNA was quantified with a NanoDrop ND-1000 spectrophotometer (Thermo Fisher Scientific), treated with DNase I (Thermo Fisher Scientific) and subjected to reverse transcription using a mix of random hexamers and oligo-dT primers and a Revert Aid first Strand cDNA Synthesis Kit (Thermo Fisher Scientific). Gene expression analysis via RT-qPCR was performed using a LightCycler480 instrument and LightCycler480 SYBR Green Master Kit reagents (Roche). Each sample was tested with four technical replicates and two dilutions. The relative *ANN5* expression level was normalized to the expression of the reference genes for vegetative and generative plant tissues (*UBC21, YLS8, RCE1, UBC9*) ^25,58^ and quantified by the ΔCt method.

#### Northern blotting

Total RNA samples were analysed on a 1.1% agarose gel and subsequently transferred to a Hybond N+ membrane by capillary transfer. To target mature and precursor rRNAs, as well as U2 snoRNA for loading normalization, oligonucleotides were radiolabeled at their 5I ends. Labeling was carried out with T4 Polynucleotide Kinase (Thermo Fisher Scientific) and γ-^32^P (SRP-201, Hartmann Analytic) according to standard manufacturer instructions. Signal detection and quantification were carried out using a Typhoon FLA 9000 Gel Imaging Scanner (GE Healthcare) and ImageQuant software (Molecular Dynamics). Oligonucleotides used for northern hybridization are listed in Supplementary Table 1.

#### Polysome profiling

Cytosolic extracts from seven-day-old seedlings and hydrated pollen grains were isolated according to previously described procedure ^59^. The polysome extraction buffer consisted of 160 mM Tris-HCl (pH 8.4), 80 mM KCl, 40 mM MgCl₂, 5.36 mM EGTA, 0.5% (v/v) Triton X-100, 50 µg/mL cycloheximide, 50 µg/mL chloramphenicol, and 0.01% (v/v) 2-mercaptoethanol. For tissue disruption, freshly collected seedlings (550 mg) were flash-frozen in liquid nitrogen and ground using a mortar and pestle. Pollen samples were ground in precooled 1.5 mL tubes using polypropylene tissue grinders with the addition of aluminum oxide. Following grinding, 600 µL or 300 µL of polysome buffer was added to the seedling and pollen lysates, respectively. The mixtures were clarified by centrifugation (twice at 16,000 × g for 15 min at 4°C). The resulting supernatants were layered onto a 10–40% linear sucrose gradient prepared in polysome buffer and separated by ultracentrifugation at 28,000 rpm for 5 h at 4°C using a TH-641 swinging-bucket rotor (Thermo Fisher Scientific). Fractions were collected using a BioComp Fractionator equipped with a UV absorbance detector. Protein-containing fractions corresponding to specific translation profiles were subsequently recovered for Western blot analysis.

### SDS-PAGE and immunoblotting analyses

For Western blot analysis, the 500 μl sucrose fractions were precipitated with an equal volume of 20% TCA/acetone (1:1 v/v) for 20 min at 4 °C. The precipitate was centrifuged at 16,000 × g for 15 min. The supernatant was removed, and the pellet was washed with ice-cold 100% acetone by inverting the tube, then centrifuged at 16,000 × g for 5 min. After removing the supernatant, the protein pellets were air-dried for 10 min, resuspended in 1.5× LDS sample buffer (Serva), and heated at 80 °C for 5 min.

Seven-day-old seedlings grown in solid MS media (or N. benthamiana leaf tissue) were flash-frozen in liquid nitrogen and ground using a mortar and pestle. Total proteins were extracted in buffer (10ml/1g tissue) 50ImM Tris-HCl pH 8.0, 150ImM NaCl, 10% glycerol, 2 mM EDTA, 5 mM MgCl2, 1% Triton X-100, 10ImM dithiothreitol, 1% protease inhibitor cocktail (Roche), for 30 min at 4 °C. Cell debris was removed via centrifugation at 16,000 × g for 15 min. For immunoprecipitation, GFP-TRAP (ChromoTek) were incubated with extracts for 1Ih at 4I°C and washed three times in wash buffer (50 mM Tris-HCl pH 8.0, 150ImM NaCl, 10% glycerol, 1% protease inhibitor cocktail before adding LDS sample buffer (4x) (Serva) and incubating for 10 min at 90 °C. Detection was carried out by SDS-PAGE and western blots using α-GFP, α-RFP (ab290, ab183628 Abcam), and α-RPS6 (PHY2025A, Phytoab) antibodies. Goat anti-rabbit-HRP (AS09602, Agrisera) was used to detect primary antibodies. Signal detection was performed using Pierce™ ECL Western Blotting Substrate (32106, Thermo Fisher Scientific). Chemiluminescence was imaged using Azure 400 AZI400-01 (Azure Biosystems, USA). Membrane loading controls were visualized by staining with Ponceau S (Sigma-Aldrich).

### Mass spectrometry analysis

For mass spectrometry analysis, cysteines were reduced and alkylated with TCEP and MMTS or DTT and IAA, and samples were digested with trypsin overnight. Resulting peptide mixtures were analyzed with LC-MS system comprised of Evosep One (Evosep Biosystems) coupled with Orbitrap Exploris 480 (Thermo Fisher Scientific) or a nanoAcquity UPLC chromatograph (Waters) with Orbitrap Velos (Thermo Fisher Scientific) or Q Exactive (Thermo Fisher Scientific) mass spectrometer. Obtained data were pre-processed with Mascot Distiller (Matrix Science) and protein identification was performed with Mascot Server (Matrix Science) using TAIR 10 database (35386 sequences). Mascot parameters included fixed cysteine modifications (Carbamidomethyl or Methylthio) and variable Oxidation on methionine, as well as trypsin digestion with 1 missed cleavage and results of the offline re-calibration with regard to precursor and fragment mass tolerance. FDR was set to 1%.

### Statistics and reproducibility

The mass spectrometry experiments were performed in biological duplicates. Microscopic observations using a confocal microscope or fluorescence stereomicroscope were conducted in three independent experiments of each type, and data from one replicate are presented in this manuscript. Statistical analyses were performed in the R environment using RStudio (ver 2025.05.1+513) and the tidyverse, ggpubr, rstatix and dplyr packages. Details of particular analyses are provided in figures’ captions.

## Acknowledgments

The research was supported by an OPUS 22 (2021/43/B/NZ3/02861) grant from the National Science Center, Poland, awarded to M.L.

## Author contributions

M.L. acquired funding, designed and performed experiments and wrote the manuscript with input from all other authors; critical manuscript review and editing were performed by MK; A.F., O.I., M.Z-P, A.G-M., N.L., I.B-F. conducted experiments, M.G., M.G-B., S.K., A.A-M. performed data analysis; A.L.S, J.H, M.K provided intellectual support.

## Ethics declarations

The authors declare no competing interests.

## Extended Data

**Extended Data Fig. 1. Analysis of POPO-1 incorporation by fixed mature pollen with modified ANN5 levels.** Confocal images of pollen grains of the wild-type Arabidopsis (Col-0), ANN5-RNAi knockdowns: KD15, KD21, overexpressing ANN5-YFP line 4. Images were collected after 60 min incubation in 0.5 µM POPO-1 in PBS. Yellow arrows depict an example of a single pollen grain that did not incorporate the dye. The dye-negative population in each line examined is attributed to technical limitations of the staining procedure of pollen with dimeric cyanine dyes rather than biological variation. Scale bar = 20 µm.

**Extended Data Fig. 2. Nucleolar dominance of ANN5 in root tip cells in Arabidopsis with ectopic ANN5 expression.** We investigated areas of cell elongation, marked by yellow squares on fluorescent stereomicroscope images, using confocal microscopy. Scale bar = 20 µm.

**Extended Data Fig. 3. Control analysis for click chemistry-based assays and Syto RNASelect staining in Arabidopsis (Col-0) seedlings. a.** Fixed root tissues incubated without 5-ethynyl uridine(–EU control) exhibit noticeable background fluorescence due to the non-specific binding of Alexa Fluor 488 azide. Incorporation of 3% bovine serum albumin (BSA) into both the blocking and washing buffers significantly mitigates this artifact. Fluorescence thresholds were determined independently for each biological replicate and applied uniformly across all control and EU-treated groups. Residual background intensity was subsequently quantified and subtracted from the corresponding EU-treated samples. Representative images demonstrate that the residual background signal does not correlate with ANN5 expression levels. Scale bar = 200 µm. b. Fluorescent stereomicroscope images show the RNA specificity of the Syto RNASelect dye; the signal is completely abolished after 30 minutes of 1 mg/ml RNase A treatment in wild-type plants but persists in ANN5–OE lines. Incomplete RNA digestion was applied intentionally to demonstrate both the specificity of the staining and the remaining excess of RNA in overexpression (OE) lines compared to wild-type (WT). Note the persistence of the signal in OE tissues under limited enzymatic treatment, confirms RNA overabundance. The lower panel shows dye-negative control root images exhibiting weak autofluorescence signal. This signal was not subtracted from dye-positive tissues, as the fluorescence threshold was set above the autofluorescence level. Scale bar = 50 µm. c. Fluorescent stereomicroscope images show validation of L-azidohomoalanine (AHA) incorporation in wild-type seedlings. Seedlings pre-incubated with the translational inhibitor cycloheximide (CHX) 50 µg/ml for 60 min exhibit a significantly reduced fluorescence signal, confirming the specificity of AHA incorporation. In contrast, translationally active control seedlings (without CHX treatment) show robust AHA incorporation, yielding intense Alexa Fluor 488 alkyne fluorescence. Optimized image acquisition parameters successfully minimized the non-specific fluorophore background signal. Because the experimental fluorescence threshold was established strictly above this baseline background level in all tested lines, no digital background subtraction was performed on the AHA-positive samples. Scale bar = 200 µm.

**Extended Data Fig. 4. 18S and 28S rRNA levels are significantly higher in seedlings with ectopic ANN5 expression.** Total RNA isolated from 20 seedlings of wild-type and ANN5-OE lines was analyzed with pulse field capillary electrophoresis. Gel image (a) and graphs (b) illustrate the difference in RNA levels between wild-type and ANN5-OE lines, generated by ProSize 3.0 software. c. The table contains measurements and calculations performed for the analyzed RNA samples. The amount of ’other RNAs’ (comprising mRNA and small RNAs) was calculated by subtracting the rRNA (18S and 28S) fraction from the total RNA yield.

**Extended Data Fig. 5. Global translation is unaffected in seedlings with ectopic ANN5 expression.** Polysome profiles were measured by sucrose density gradient fractionation of total extracts from 7-day-old seedlings of wild-type and ANN5-OE2. Plots show overlaid polysome profiles of wild-type and ANN5-OE2 from the small ribosomal subunit 40S peak to the end of the polysomes. All traces were aligned with respect to the monosome peak height along the x-axis. Data from three independent experiments are shown.

## Supplementary information

**Supplementary Table 1. Oligonucleotides used in this study.**

**Supplementary Data 1. Proteomic dataset of proteins co-purified with ANN5 from hydrated pollen and 7-day-old seedlings.**

---

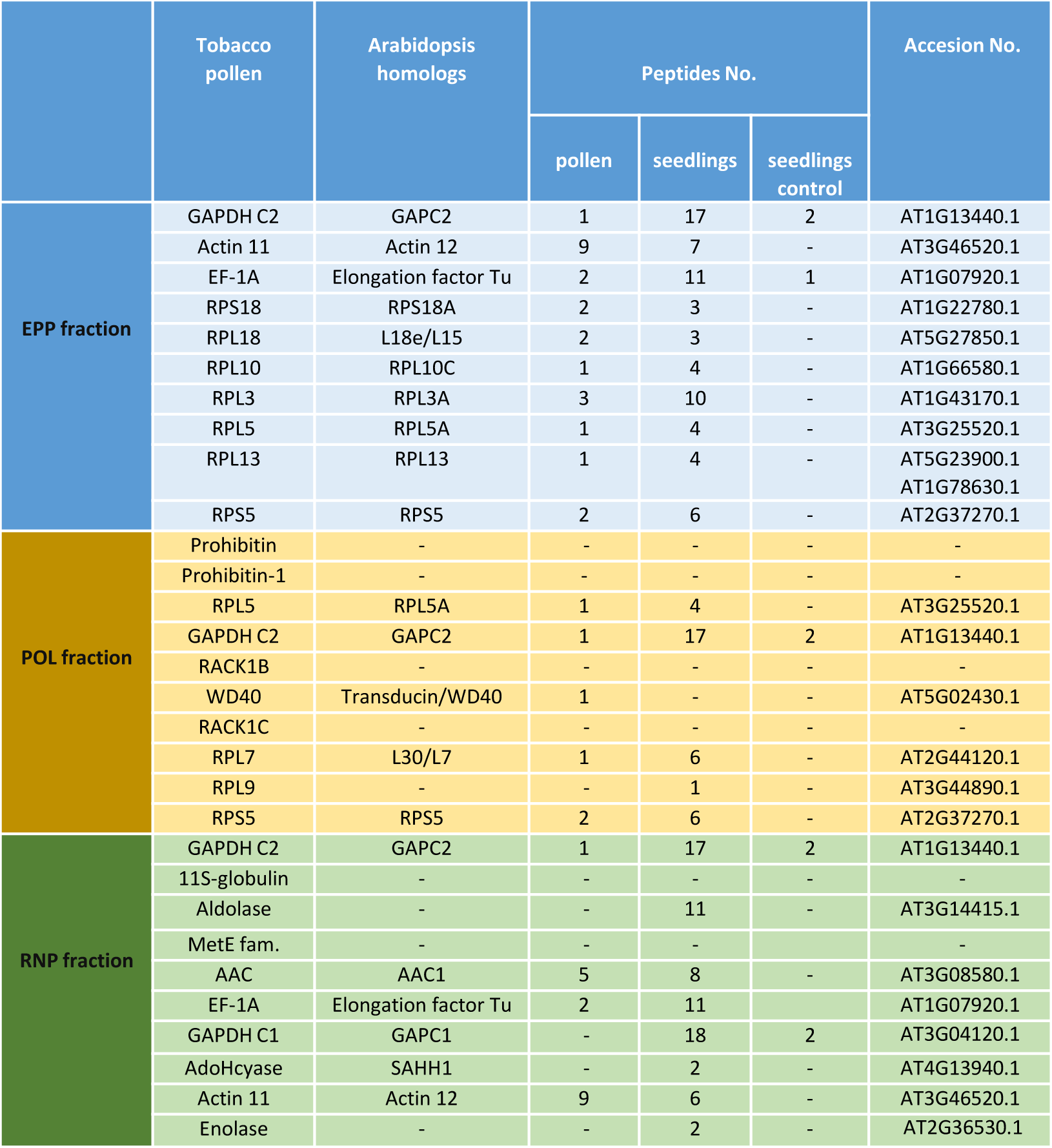

