## Supplementary Figures for "Annexin 5 is a key player in orchestrating the pollen sequestrome"

Supplementary Figure 1.

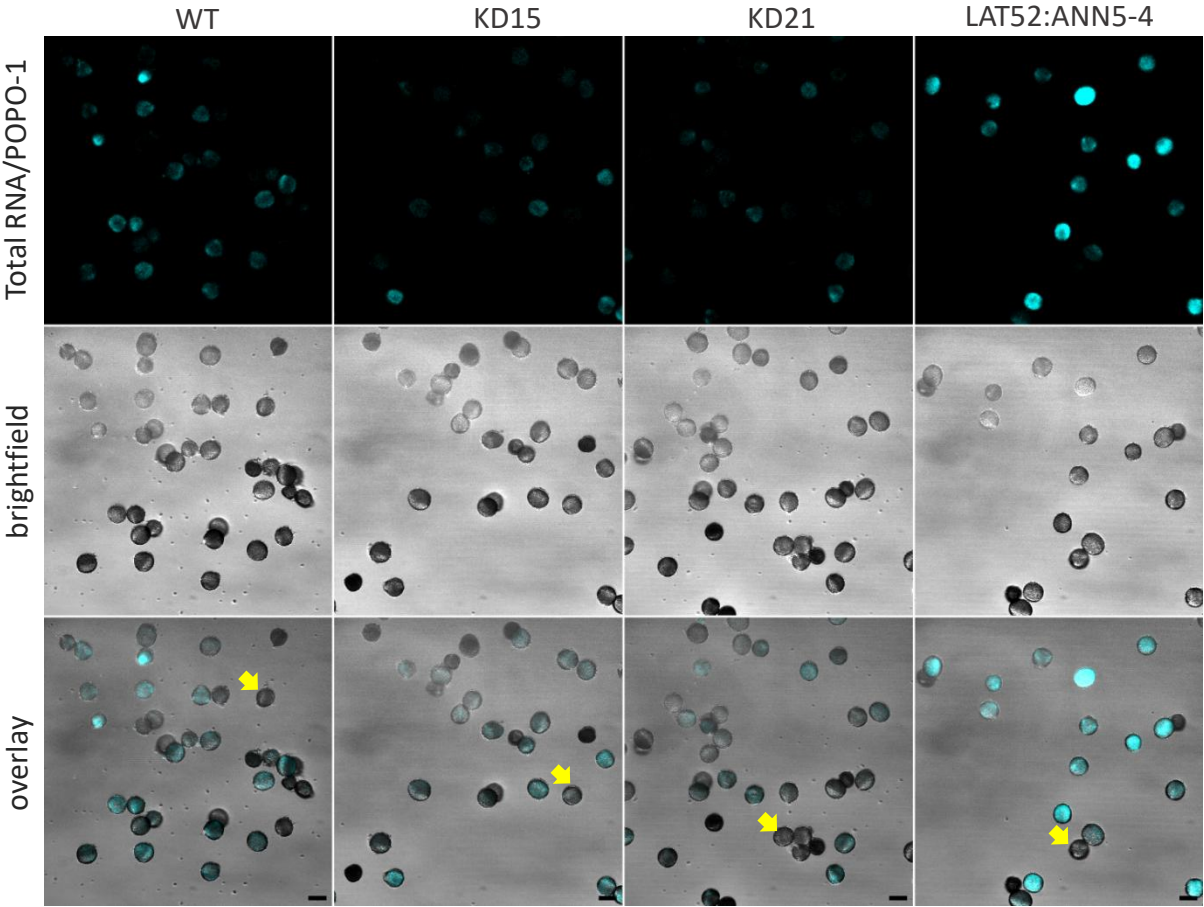

Supplementary Figure 2.

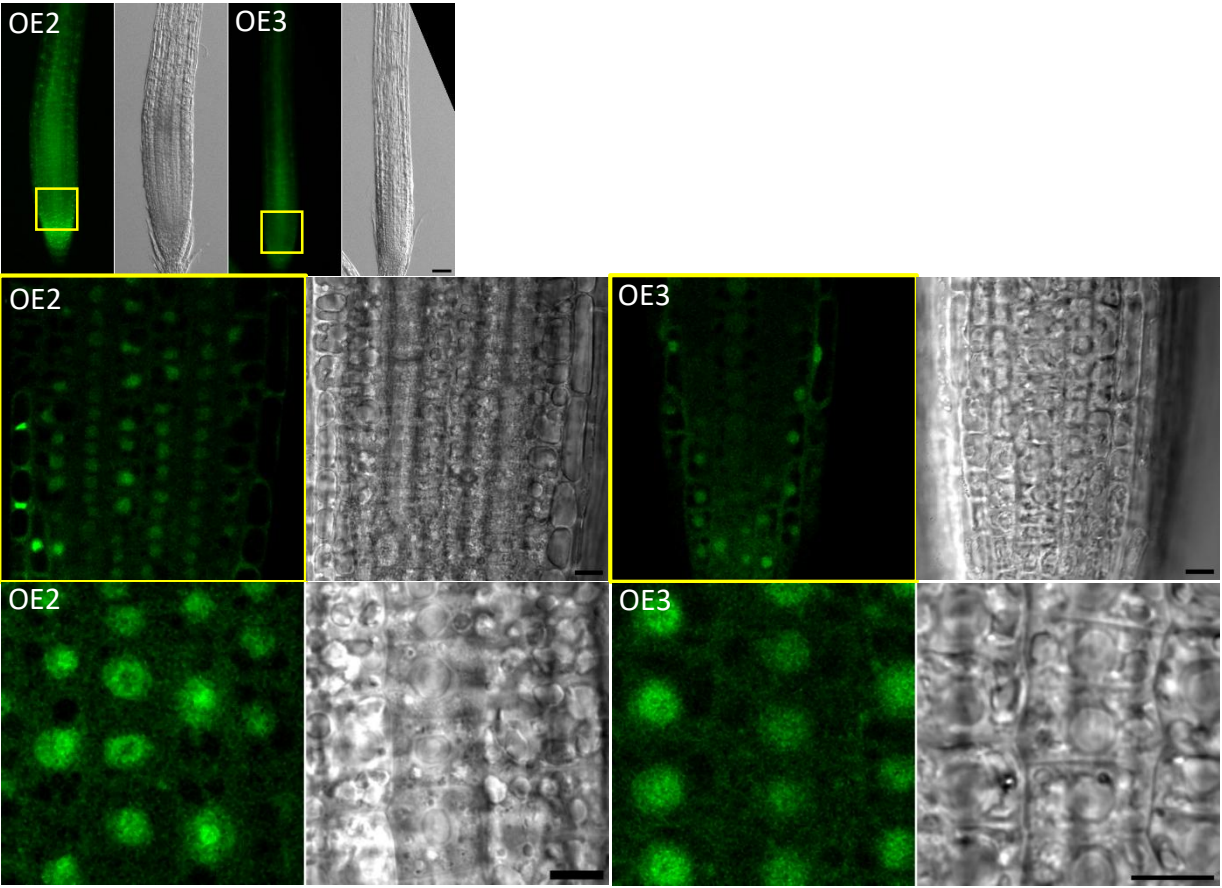

Supplementary Figure 3.

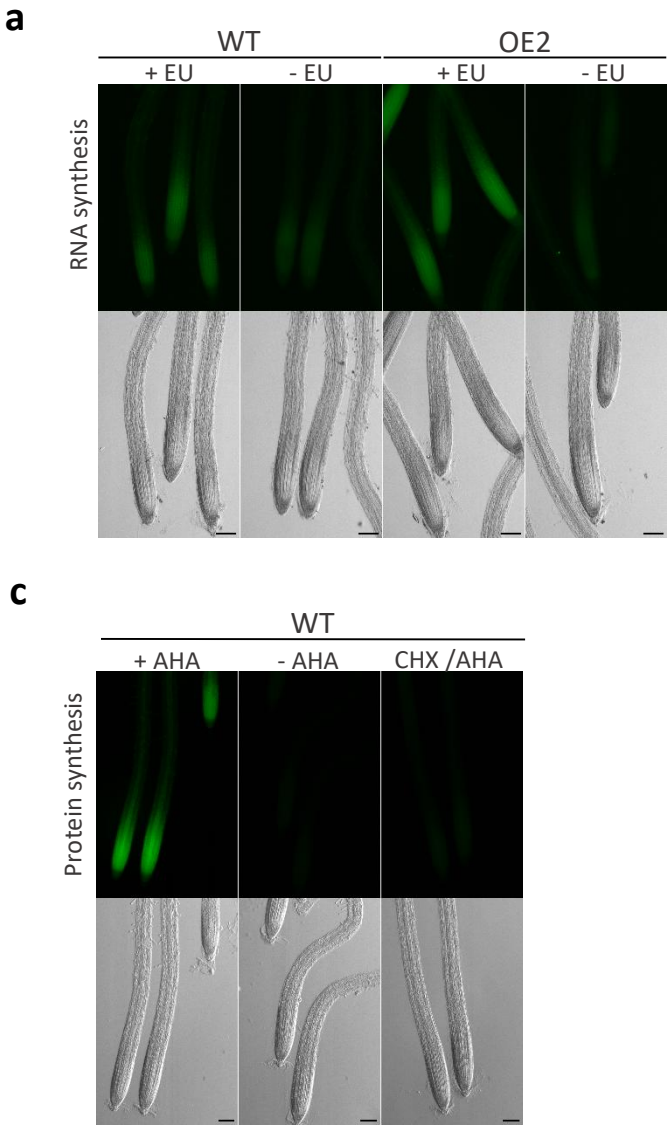

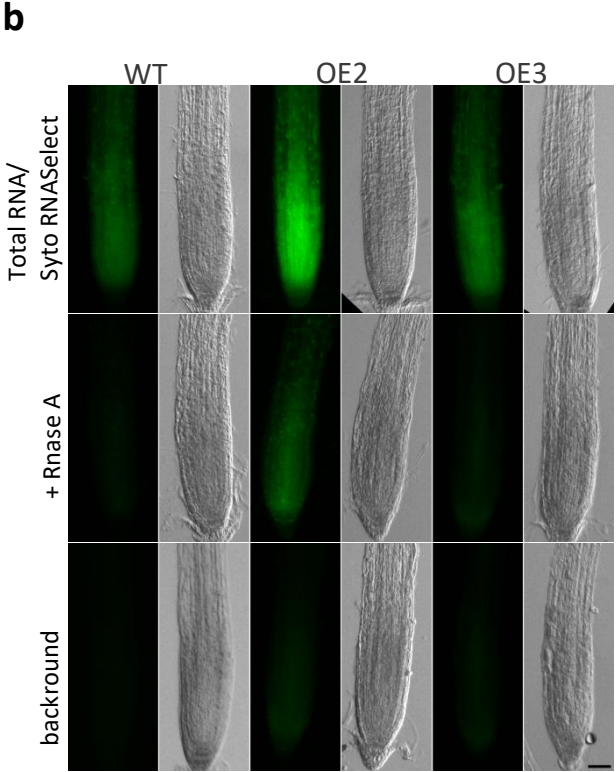

Supplementary Figure 4.

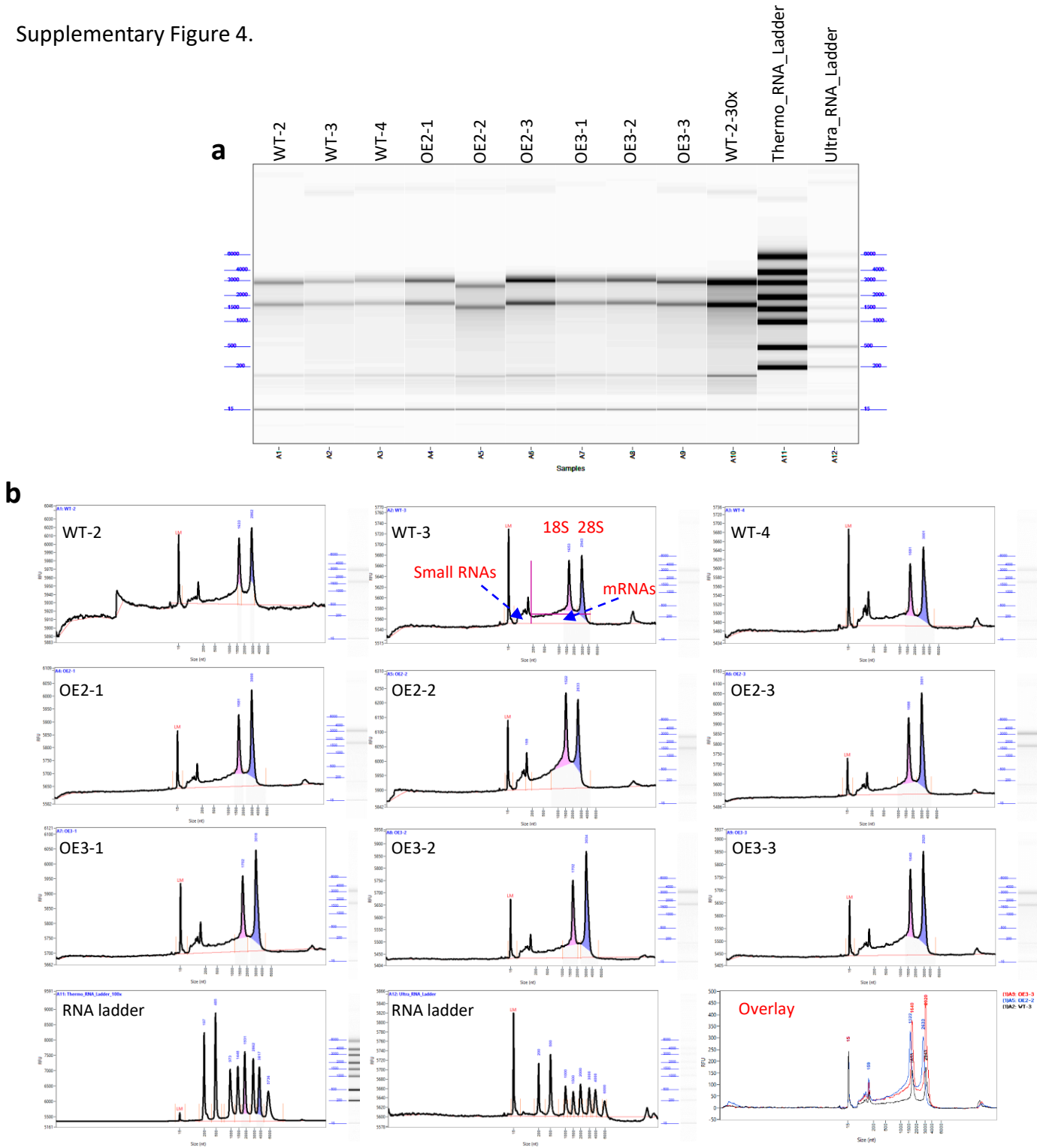

**c**

| Genotype | Total RNA<br>Qubit 3.0 | Total RNA<br>Femto Pulse | 18S | 28S | Other RNAs |
| --- | --- | --- | --- | --- | --- |
|  | [ng/μl] | [ng/μl] | [ng/μl] | [ng/μl] | [ng/μl] |
| WT-2 | 33 | 0.47 | 0.10 | 0.11 | 0.25 |
| WT-3 | 28.2 | 0.41 | 0.12 | 0.09 | 0.20 |
| WT-4 | 31.2 | 0.47 | 0.11 | 0.13 | 0.23 |
| OE2-1 | 48.6 | 0.70 | 0.17 | 0.21 | 0.32 |
| OE2-2 | 61 | 0.86 | 0.38 | 0.19 | 0.29 |
| OE2-3 | 62 | 1.14 | 0.42 | 0.37 | 0.35 |
| OE3-1 | 46 | 0.61 | 0.17 | 0.18 | 0.26 |
| OE3-2 | 47 | 0.70 | 0.22 | 0.22 | 0.27 |
| OE3-3 | 51 | 0.88 | 0.25 | 0.24 | 0.39 |

Supplementary Figure 5.

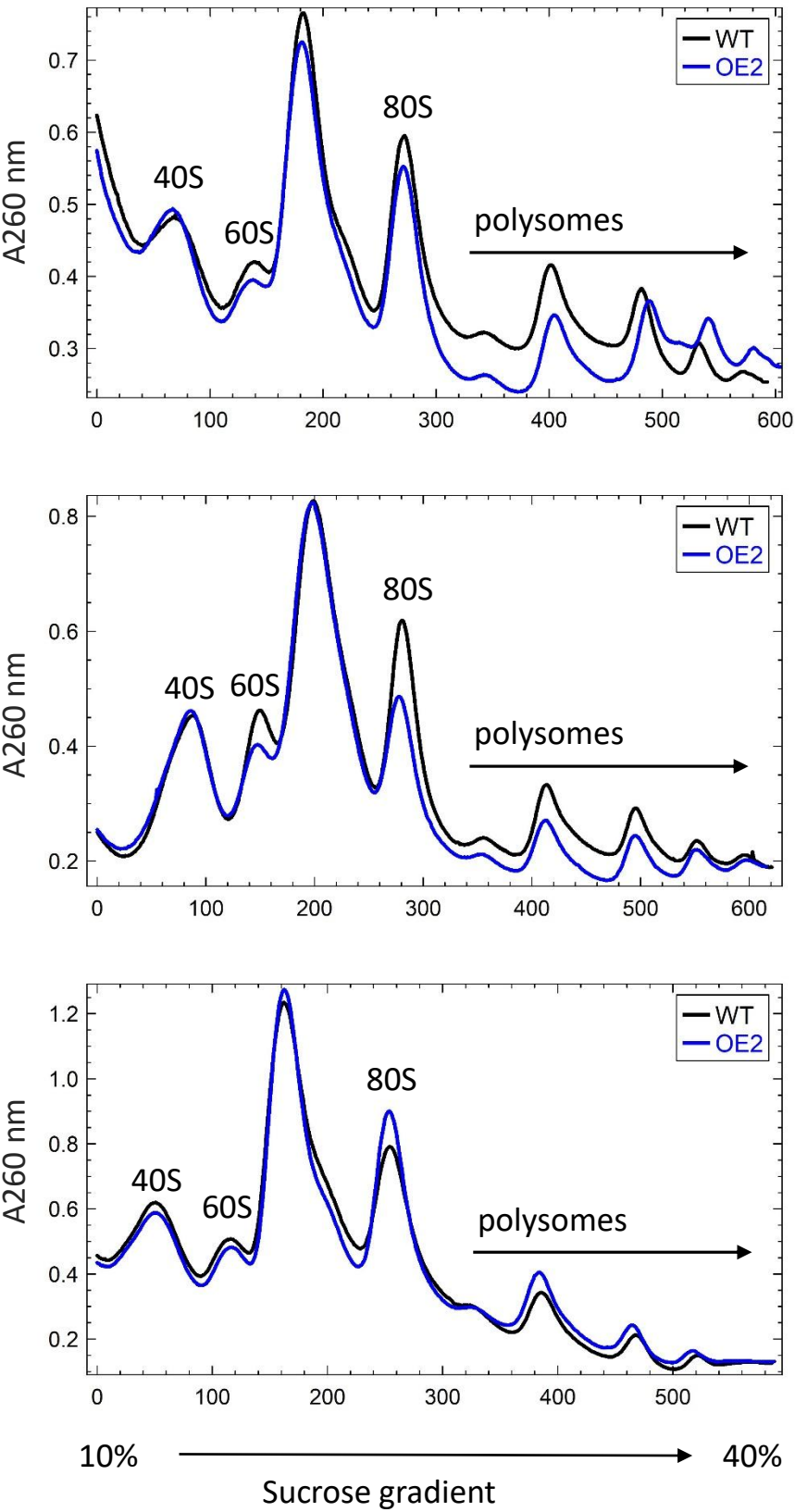
